# Initial Characterization of Canine T-cell Receptor Repertoires Using RNA-seq Data from Different Diseases and Tissues

**DOI:** 10.1101/2024.02.14.580390

**Authors:** Mengyuan Zhang, Kun-Lin Ho, Shaying Zhao

**Affiliations:** Department of Biochemistry and Molecular Biology, Institute of Bioinformatics, University of Georgia, Athens, GA 30602, USA

## Abstract

The dog serves as a key translational model in cancer immunotherapy. Understanding the T cell receptor (TCR) repertoire is needed for various cancer immunotherapies. Compared to humans where >300 million TCRs have been identified, <100 canine TCRs are reported. To address this deficiency, we assembled >200,000 complete TCR complementarity-determining region 3 (CDR3) sequences from RNA-seq data published for ∼2,000 canine samples of blood, lymph node, and other tissues, of which 613 are tumors. We collected 1,324 human RNA-seq samples to compare the similarities and differences in T-cell repertoires between humans and dogs. Notably, our analysis revealed distinct variable gene usage patterns between blood samples and solid tissues in both canine and human samples for TRA and TRB loci. Moreover, our investigation led to the discovery of novel V gene and allele candidates in the canine genome. Our findings also revealed that the canine CDR3 resembled human CDR3 in terms of length and motifs. Additionally, our study unveiled shared traits in cancer TCRs between dogs and humans, including longer lengths and higher hydrophobicity of private CDR3s. Our results indicated the diversity of canine to be more comparable to that of humans than mice. Our study provides an initial landscape of the canine TCR repertoire, highlighting both its similarities and differences with the human counterpart, thus laying the groundwork for future research in comparative immunology and vaccine development.

## Introduction

An individual has up to a billion of unique T-cell clonotypes^1,2^, each identified by its TCR. A TCR is a heterodimer, consisting of α and β chains in ∼95% T cells, and δ and γ chains in ∼5% T cells. CD8 and CD4 T cells, activated by major histocompatibility complex (MHC) -I and MHC-II presented antigens respectively, and which mediated tumor-cell killing, have αβ TCRs^3,4,5^. The TCR repertoire, consisting of TCRs of all T cells, harbor the information of the history of antigen-exposure in an individual. Thus, TCR repertoire is a key element in cancer immunotherapy development.

Canine cancers more faithfully represent their human counterparts, enhancing their translational values. Unlike rodent models, canine cancers arise naturally in animals with intact immune systems, thereby recapturing the essence of human cancer. As companion animals, dogs share the same environments as humans and are exposed to many of the same carcinogens^6,7,8,9^. Indeed, environmental toxins, advancing age, and obesity are known risk factors for canine cancer^10^. Studies by us^11^ and others^12,13^, show a strong dog-human molecular homology for various cancer types/subtypes.

While >300 million human TCRs are defined^14^, <100 canine TCRs are published^15^. The canine TCR repertoires remain largely uncharacterized, preventing the effective use of the canine model in immune research. To address this deficiency, we take advantage of the RNA-seq published for ∼2,000 canine samples and conduct the study below.

## Results

### Sample collection and QC

We collect paired-end RNA-seq data of 1,361canine samples of 1,063 dogs from the SRA database^16^. These include specimens from blood and lymphocyte where the T cells are circulating, as well as samples from mammary, colon, skin and other tissues harboring T cells that are resident.

We then performed comprehensive QC analysis as described^11,17^, by investigating sequencing amount and quality, reference-genome mapping properties, tumor-normal sample pairing accuracy, as well as canine breed validation and prediction. A total of 1,293 samples from 976 dogs passed our QC measures. In addition, a total 326 dogs have their breed validated or predicted.

### TRUST4 and MiXCR comparison

TRUST4^18^ and MiXCR^19^ serve as widely used TCR reconstruction tools. TRUST4 is designed to reconstruct immune receptor repertoire for bulk and single-cell RNA-seq data, whereas MiXCR is a versatile tool suitable for various sequencing data types. To employ the optimal tool and obtain accurate CDR3, we applied both two tools to the entire RNA-seq data. We compared the results obtained from TRUST4 and MiXCR in two aspects: quantity and quality. Before analysis, we extracted complete CDR3 based on amino acid sequences which start with conserved cystine(C) and end with Phenylalanine (F)^15^. In terms of quantity, while there was significant overlap between the complete CDR3s identified by MiXCR and TRUST4 (Fig.1A), TRUST4 identified more non-overlapping CDR3s compared to MiXCR. Regarding quality, we examined the length distribution (Fig.1B) and motif conservations using shuffling strategy (Fig.1C). Utilizing the overlap CDR3s as a reference, we assessed differences in length distribution and motif conservation between shared CDR3s and unique CDR3s identified by TRUST4 and MiXCR separately, using JSD and PCC metrics. The results indicate the CDR3s gained from TRUST4 are comparable to CDR3s obtained by MiXCR in quality. While previous studies reported that TRUST4 has a higher false positive rate. We conducted further examination on the TCR contigs assembled by both software corresponding to the complete CDR3. We apply blast^20^ to align the TCR contigs to reference TCR genes. The alignment length of TCR contigs assembled by TRUST4 is longer than that of MiXCR (Fig.1D). The identity score of TCR contigs from TRUST4 to the references is similar to the that of MiXCR. Based on these analyses, we have chosen to utilize the CDR3 and TCR contigs reported by TRUST4 for downstream analysis.

**Figure 1.**
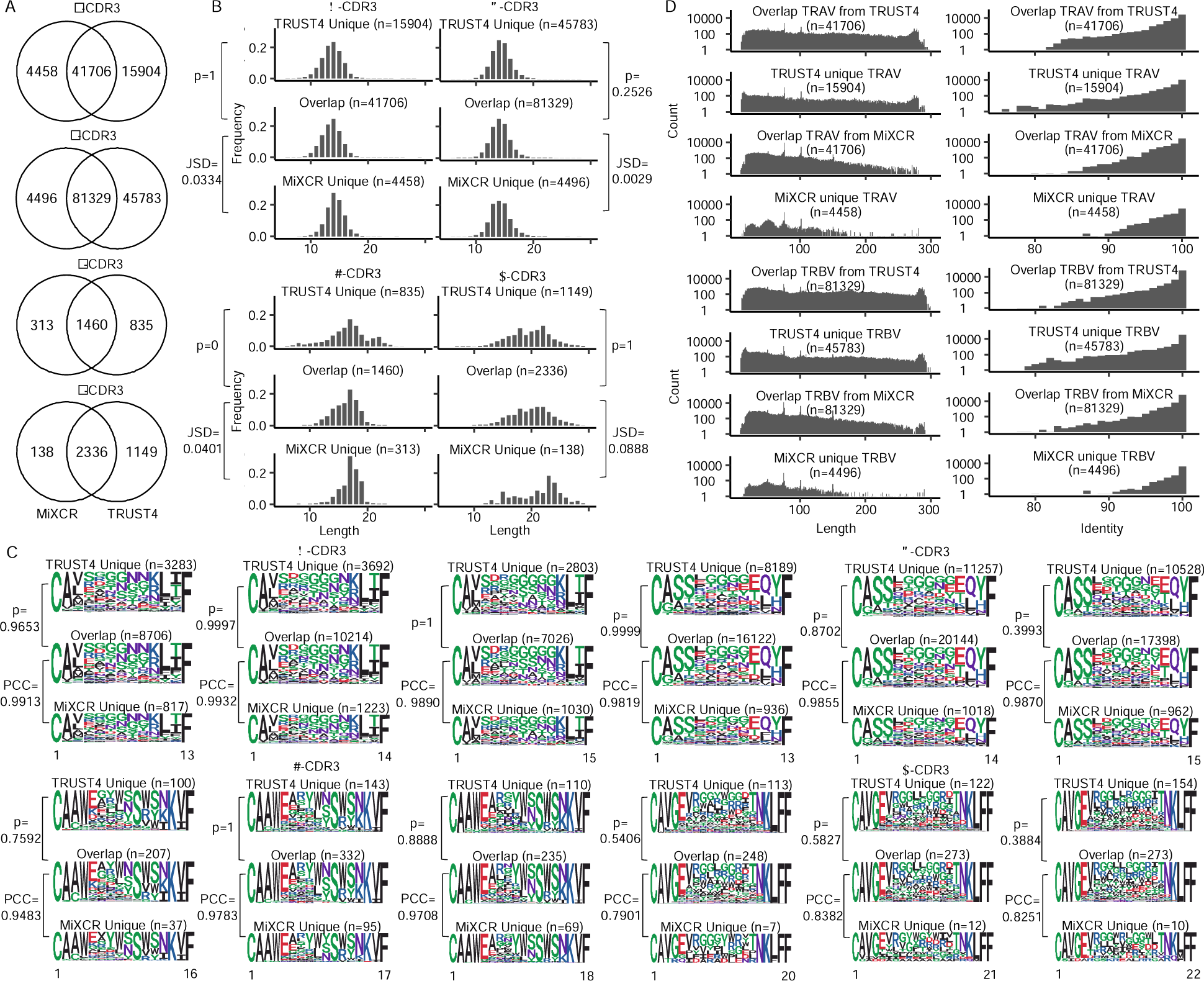
TRUST4 identifies more TCRs than MiXCR with canine RNA-seq. A. Venn diagrams indicate the total number numbers of complete CDR3s identified by both software and by only one software. B. Length distribution of shared and software-unique complete CDR3s. p-values, obtained via shuffling analysis (see Methods), indicate no significant difference between the distributions compared. JSD: Jensen–Shannon divergence. C. Motifs of amino acid sequences of shared and software-unique complete CDR3s. p-values, obtained via shuffling analysis (see Methods), indicate no significant difference between motifs compared. PCC: Pearson coefficient correlation. D. The length distribution and identity of TCR contigs of TRUST4 unique, overlap, and MiXCR unique.

### TRAV and TRBV gene usage

Gene usage analysis is fundamental for deciphering the mechanisms underlying TCR diversity, primarily driven by V(D)J recombination. The CDR1, CDR2, and partial CDR3 encoded by variable genes play a vital role in interacting with MHC and peptides. To obtain a profound understanding of V(D)J recombination, we investigated the human and dog TRAV and TRBV gene usage across various tissues. Our findings revealed that the gene usage profiles of TRAV and TRBV in human blood are different from that from mammary tissues (Fig.2A). Similarly, distinct gene usages are observed between canine blood and solid tissues. Interestingly, we observed the divergent TRAV and TRBV gene usage between lymphoid tissues and blood, while noting similarities to solid tissues (Fig.2B). We observed a cross-species conservation in the predominant TRBV gene usage that TRBV20 exhibiting the highest frequency in both human and canine samples. This analysis highlights that the gene use profiles of humans and dogs are similar.

**Figure 2.**
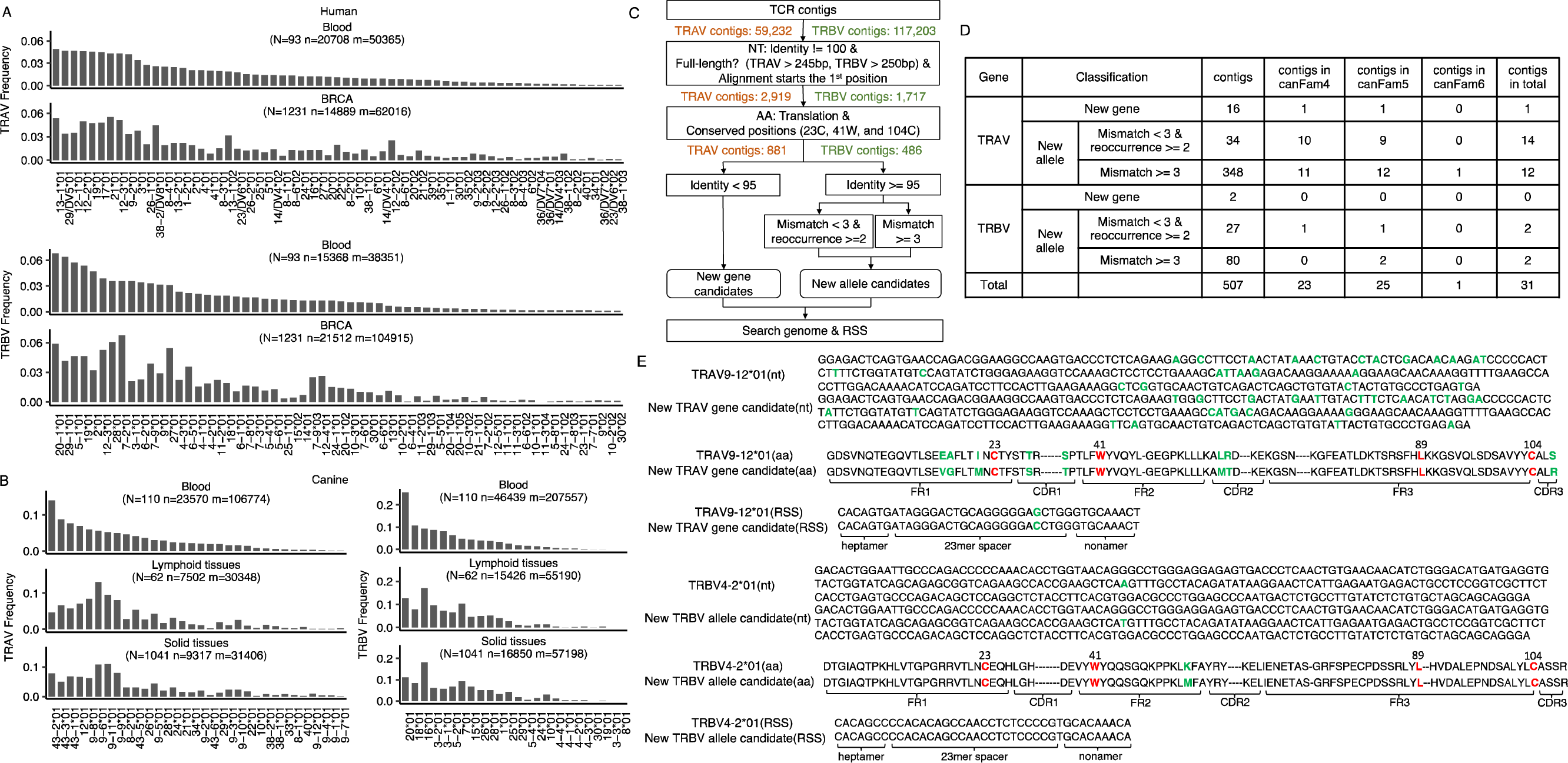
Analysis of Gene usage and discovery of new genes and alleles. A. Distribution of TRAV and TRBV gene usage in human blood and breast cancer samples. N represents the number of samples, n represents the number of CDR3 sequences, while m denotes the number of supporting reads for each CDR3. B. Gene usage analysis of TRAV and TRBV in dog blood, lymphoid tissues, and solid tissue samples. C. A flowchart of pipeline of new canine TRAV and TRBV genes and alleles identification. D. Summary table of new canine TRAV and TRBV gene and allele candidates mapped to genomes. E. Alignment results showcasing examples of new canine TRAV gene candidate and new canine TRBV allele candidate compared to known references.

### Exploration of new variable genes and alleles in canine

In comparison to the 77 known TRBV genes in humans, only 36 TRBV genes have been identified in canines^15^. Additionally, while humans exhibit multiple alleles for each V gene, only one allele per V gene has been characterized in dogs. To enhance the TCR germline repertoires in canines, we implemented a rigorous analysis to uncover new V genes and alleles in dogs (Fig 2C). Following the exclusion of low-quality contigs, we identified 507 new gene and allele candidates (Fig.2D). Subsequently, we searched contigs in three genomes (canFam4, canFam5, and canFam5) and recombination signal sequences (RSS). As a result, we discovered one new TRAV gene candidate and 26 new TRAV allele candidates, along with 4 new TRBV allele candidates (Fig.2D). Experimental validation of these candidates is forthcoming. Moreover, we presented alignment results showing the comparison of a TRAV new gene candidate and TRBV new allele candidates with a reference at nucleotide and amino acid levels, and RSS (Fig.2E). This endeavor contributes to improving our understanding of canine TCR germline repertoires.

### The length distribution and motif conservation of CDR3

The length distribution and motif conservation of CDR3 play pivotal roles in TCR diversity and functionality. As the most diverse region of TCR, CDR3 directly interacts with MHC molecules and peptides, thereby influencing immune responses. We examined the length distribution of four types of CDR3 among distinct tissues in humans and dogs. The results indicate the length range of α and β-CDR3 is more constrained compared to γ and δ-CDR3 in dogs (Fig.3A), agreeing with findings in humans. Notably, we observed that the length of γ and δ-CDR3 in dogs tends to be longer than in humans, with a wider length range. The motifs of CDR3 contribute to the binding specificity and affinity of interactions with pMHC. We investigated the motifs of dominant lengths of α and β-CDR3 in distinct human and dog tissues (Fig.3B). Our results revealed highly conserved motifs across canine tissues, resembling the motif patterns observed in human CDR3. This analysis suggests similarities in immune response mechanisms between humans and dogs.

**Figure 3.**
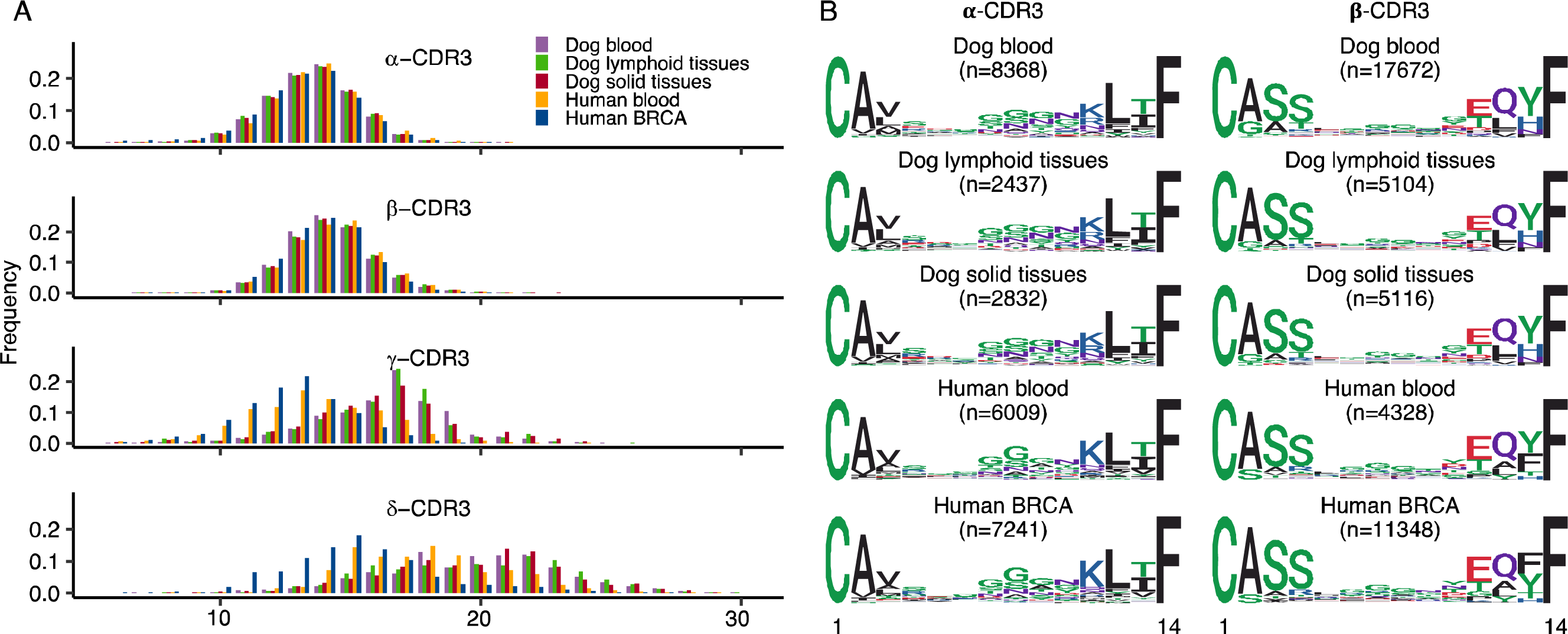
The length distribution and sequence motifs of canine α- and β-CDR3 resemble those of their human counterparts. A. Histograms illustrating the length distributions of complete CDR3 amino acid sequences in dog blood, dog lymphoid tissues, dog solid tissues, human blood, and human breast cancer samples. The x-axis represents the length range from 6 to 30. B. Amino acid conservation of α- and β-CDR3 across dogs and humans. Analysis focused on α- and β-CDR3 sequences with 14 amino acids for this comparison.

### The clonality and diversity

The clonality and diversity of TCR determine the amount of the antigens TCR that can be recognized. We analyzed the clonality and diversity of TCR repertoires across distinct groups in humans and dogs. The results demonstrated the clonality and diversity of dog TCR repertoires resembled that of humans. We also observed that the BLO and NL samples exhibited more clonotype counts and higher read counts compared to other resident tissues. Interestingly, as depicted in Fig.4A, samples from TCL displayed distinct characteristics, with the highest clonality but the lowest diversity among the 11 canine groups. These findings align with the pathogenesis of TCL, where specific clonotypes undergo marked expansion^21^. Furthermore, to explore the relationship between MHC-I expression and TCR repertoires, we employed Kmer-based paired-end read de novo assembler and genotyper (KPR)^17^ to estimate MHC-I reads for each sample. The results indicated a robust positive correlation between MHC-I expression levels and TCR expression levels (Fig.4B).

**Figure 4.**
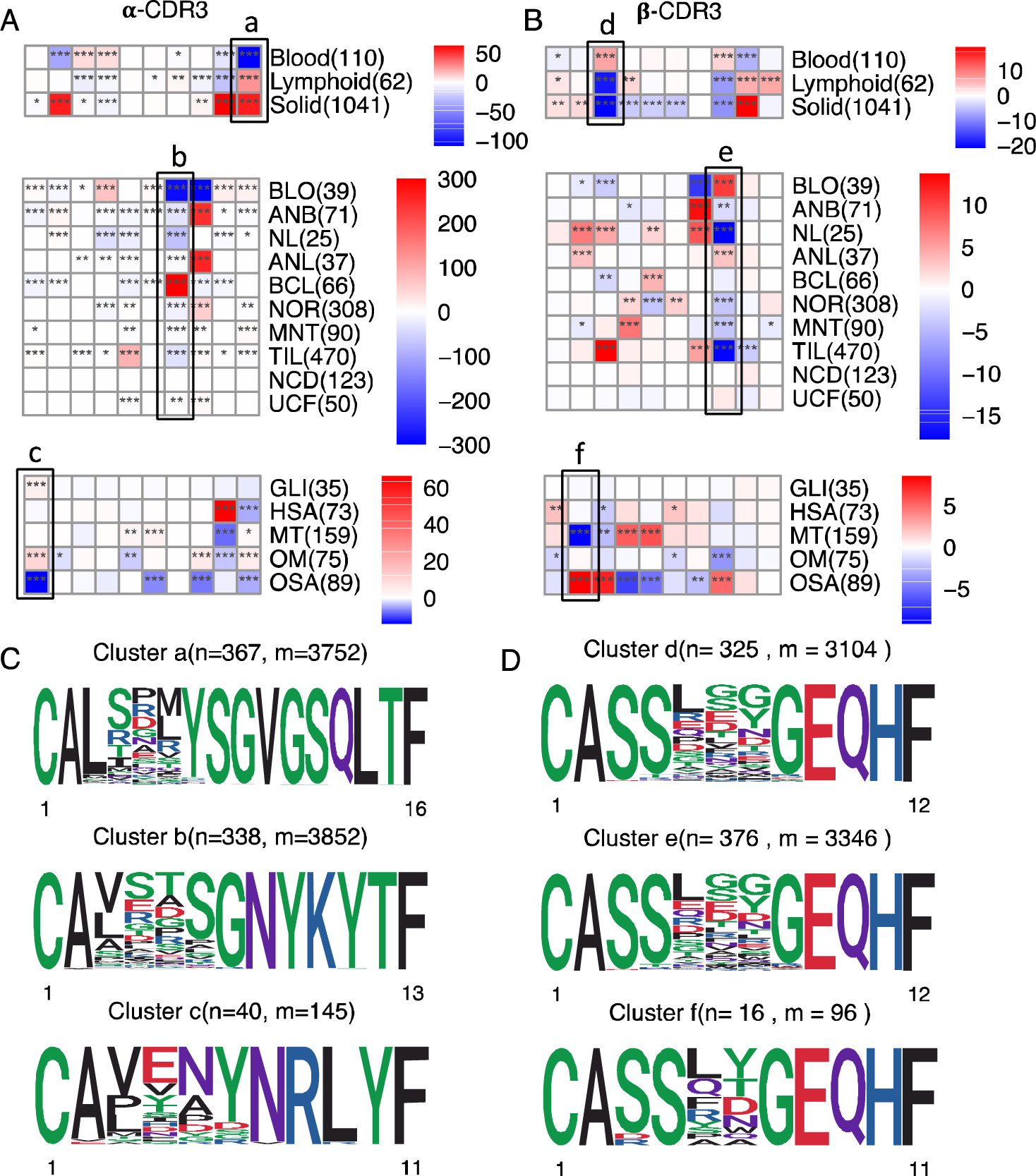
Canine α and β-CDR3 sequence clustering, enrichment and depletion analysis. A and B. Heatmaps illustrating the α and β-CDR3 enrichment and depletion across various tissues, and cancer types. Highlighted clusters denote significant clusters in the heatmap. Colors: red indicates enrichment; blue indicates depletion. (*: p-value < 0.05; **: p-value < 0.01; ***: p- value < 0.001) C and D. Representation of motifs for clusters identified in panels A and B. n represents the number of CDR3 sequences, while m denotes the number of supporting reads for each CDR3.

### Enrichment analysis

To explore the enrichment or depletion of CDR3 clusters in specific tissue types and cancers, we used clusTCR^22^, an unsupervised clustering tool for CDR3 sequences, to identify the CDR3 clusters. Subsequently, the Fisher exact test was employed to assess the enrichment and depletion. To reduce the possibility of false enrichment results due to samples with abnormally high expression of certain CDR3 sequences, we conducted QC measures and only highlighted the validated enrichment results (Fig.5). We observed α and β-CDR3 clusters exhibiting enrichment in distinct tissue types (Fig.5A). Cluster b displayed extreme enrichment in BCL samples but depletion in other groups, that potentially indicated α-CDR3s in cluster b in BCL samples are from activated T-cells. We also explored the motifs of those enriched or depleted CDR3 clusters. Intriguingly, the cluster d and e share a highly identical motif. Those highlighted β-CDR3 clusters are highly conserved at the first four and last five amino acids (Fig.5B).

**Figure 5.**
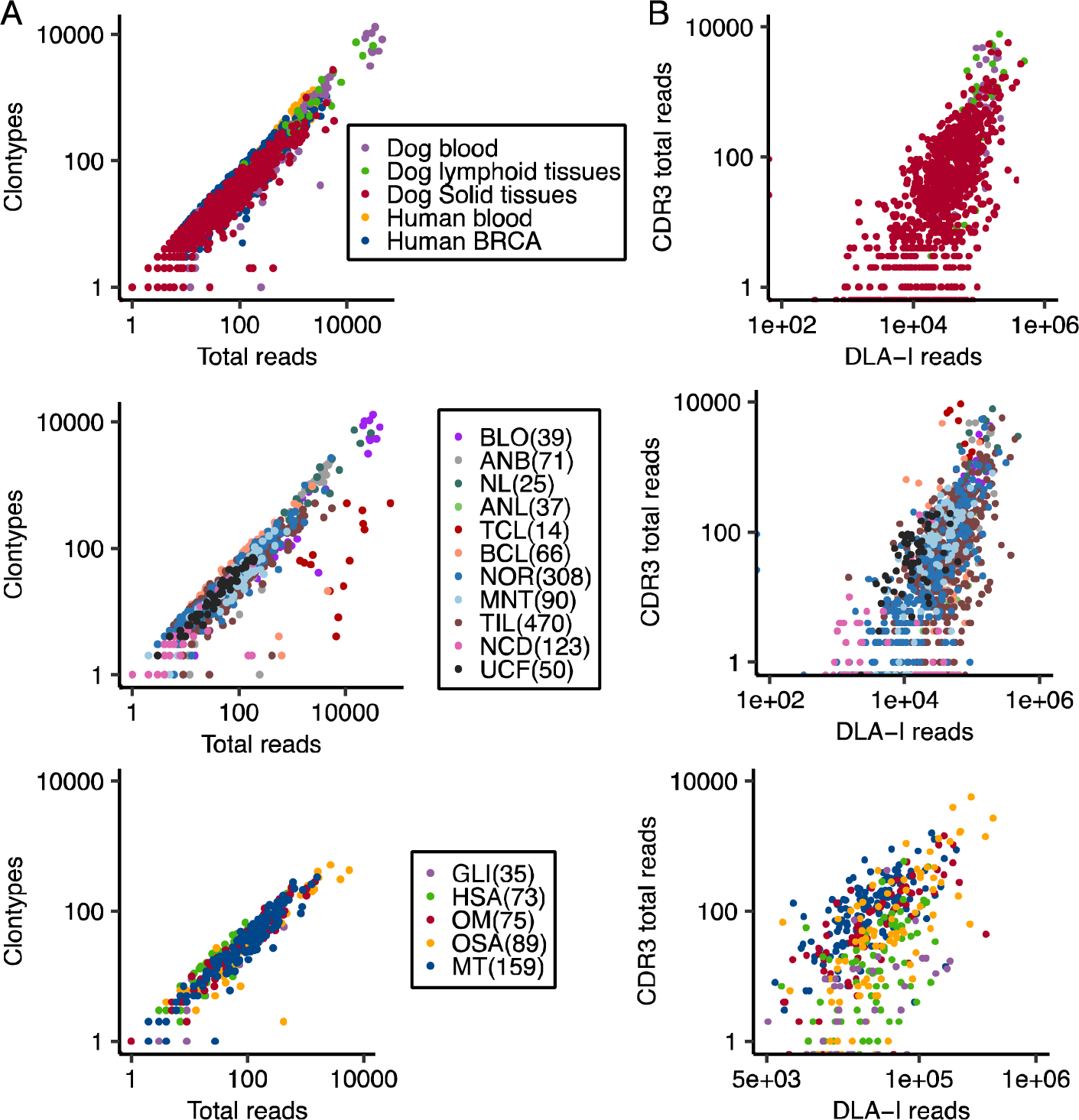
Discovery of T cell clonality and diversity. A. Plots depicting the number of clonotypes versus the total reads of CDR3 in distinct categories (top), tissues (middle), and cancer types (bottom). Different colors represent different categories, tissues, and cancer types. B. Plots illustrating the total reads of CDR3 versus the total reads of DLA-I in diverse categories (top), tissues (middle), and cancer types (bottom).

### Tumor public and private CDR3

Understanding the interplay between private and public CDR3s provides valuable insights into the diversity, specificity, and dynamics of T-cell responses across individuals. The public or private CDR3s are defined as the CDR3 being shared among more than two samples or only identified in one sample in this study. In this study, we only classified the public and private CDR3 within tumor samples. We investigated the intersection α and β-CDR3 across tumor private, tumor public, circulating and resident sets. The results (Fig.6A) shown more β-CDR3 were identified by TRUST4 compared to α-CDR3, while more α-CDR3 are shared by multiple samples. We devolve into the tumor public α and β- CDR3, a significant prevalence of α-CDR3s was observed across multiple samples (Fig.6B). The results indicated the diversity of β-CDR3 is more diverse than α-CDR3, which consist with an extra D region involved in the somatic recombination of β chain. A previous study^23^ focusing on human tumor-infiltrating T-cells noted that private β-CDR3 sequences are significantly longer than their public counterparts (Fig.6C). Furthermore, the study found that the middle three amino acids in private β-CDR3 sequences are notably more hydrophobic than those in public sequences. In our analysis of both α and β-CDR3 sequences in dogs exhibited similar length trends as observed in humans (Fig.6D). Moreover, for dog β-CDR3s, our hydrophobicity analysis yielded results consistent with those found in humans. Interestingly, the hydrophobicity profiles of α-CDR3 are completely opposite to β-CDR3 which might uncover the function of α-CDR3 is different from β-CDR3. These findings suggested that the tumor private CDR3s in canine are highly potential related to neoantigens and could be putative vaccine targets as human tumor private CDR3s.

**Figure 6.**
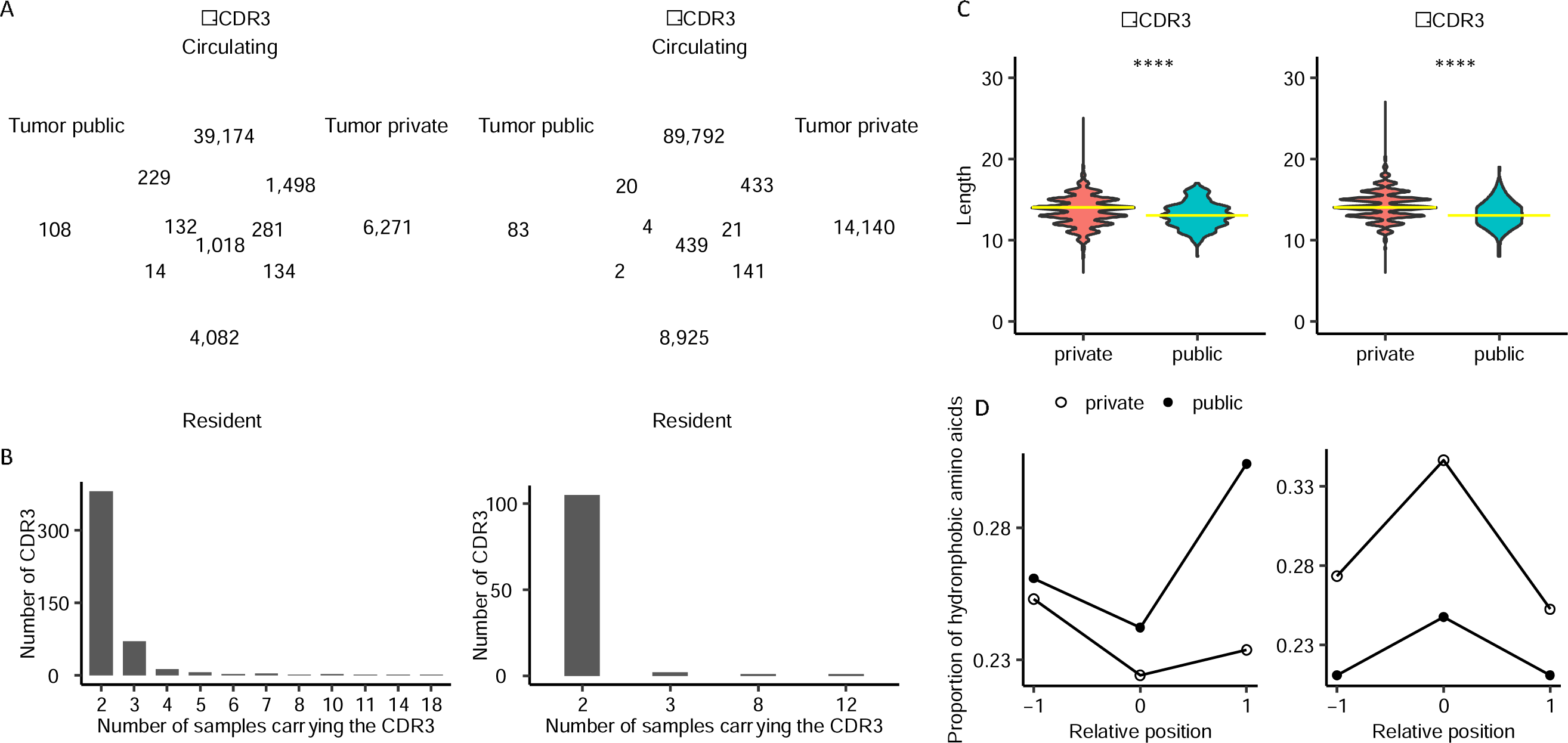
Public and private α- and β-CDR3 amino acid sequences have different lengths and proportions of hydrophobic residues. A. Venn diagram illustrating the intersection of α- and β-CDR3 sequences among T-cell infiltrating tumors (TIL), circulating samples, and resident samples. Left panel: α-CDR3; Right panel: β-CDR3. B. Frequency distribution of samples carrying TIL public α- and β-CDR3 sequences. C. Comparisons of private and public α- and β-CDR3 sequences lengths in TIL samples, respectively. Median values are indicated by yellow lines. The Wilcoxon test calculated the P- values (****: p-value < 0.0001). D. Analysis of hydrophobicity in the middle three amino acids of private and public α- and β- CDR3 sequences.

### Comparison of **β**-CDR3 across dogs, mice, and humans

Mouse has been widely used as animal model for biomedical research, drug discovery and vaccine development. A comparative analysis of mouse and canine CDR3 is essential for gaining insights into the similarities between dog and human CDR3s, and mouse and human CDR3s. To conduct this comparison, we gathered β-CDR3 sequences for these species from TCR-seq data. Using clusTCR^22^, we clustered the dog and human CDR3s, as well as the mouse and human CDR3s, separately. Interestingly, while the software could equally cluster the dog and human CDR3s, the CDR3s of mice formed more clusters compared to humans (Fig.7A, D). Furthermore, more clusters were identified in canine and human CDR3s compared to mice (Fig.7B, E). Upon summarizing the cluster size across 100 replicate analyses, we noted that the cluster size of mice was larger than that of dogs (Fig.8C, F). This suggests that the CDR3 repertoire in dogs is more diversified than that in mice, yet comparable to that in humans. Both canine CDR3s and mouse CDR3s could be clustered alongside human CDR3s, indicating a degree of similarity in CDR3 sequences among dogs, mice, and humans. This comparative analysis supports the notion that canines could serve as valuable animal models for studying immune responses and immunotherapy for humans.

**Figure 7.**
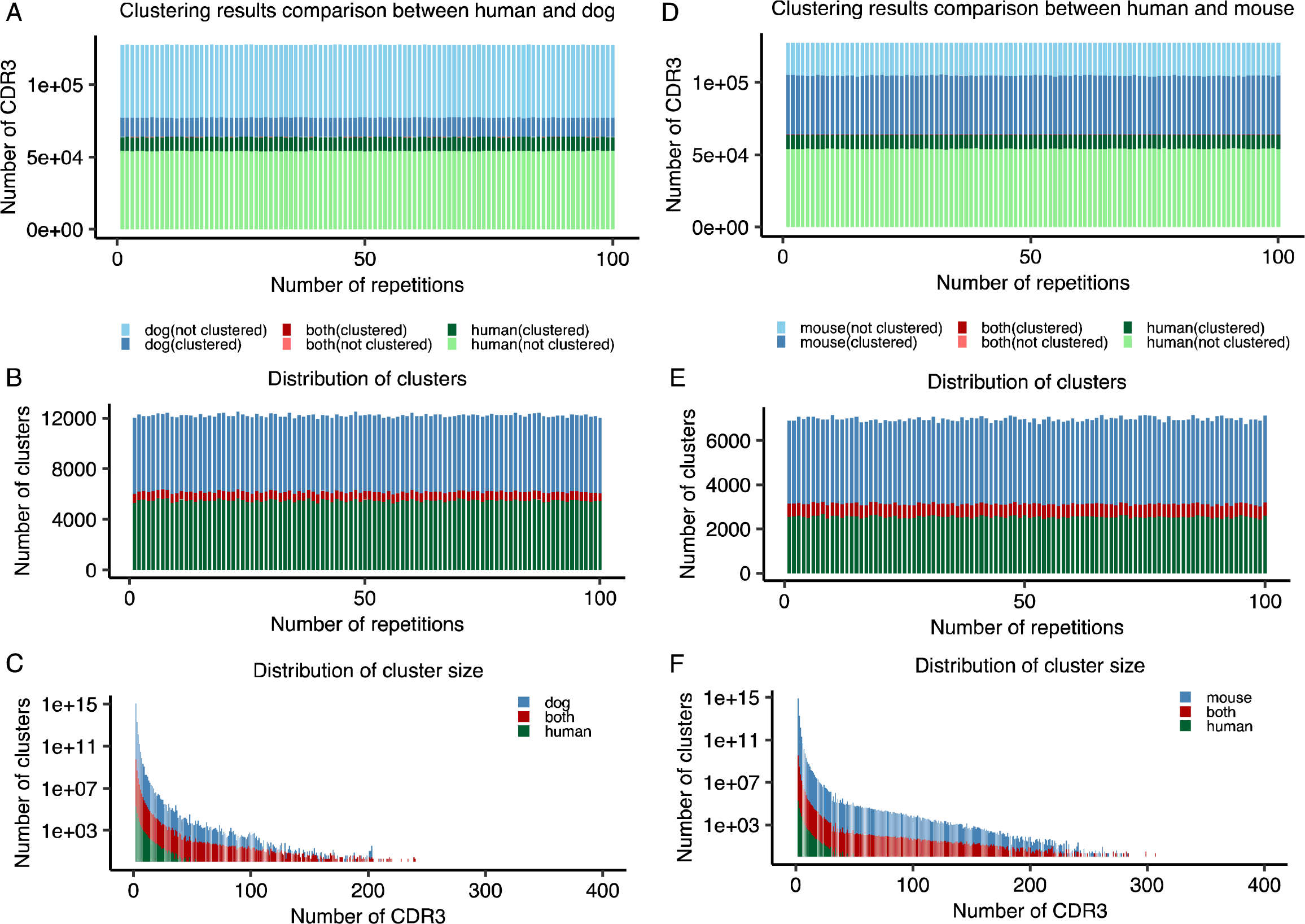
The diversity of canine β- CDR3 is more akin to that observed in human β- CDR3 than in mouse β- CDR3. A. Comparison of the number of β-CDR3 sequences clustered or unclustered from dogs and humans across 100 repetitions. Each stacked bar represents one repetition. Blue, red, and green represent the counts of β-CDR3 sequences clustered from dogs, both species, and humans, respectively. Light blue, orange, and light green bars represent the counts of unclustered β-CDR3 sequences from dogs, both species, and humans, respectively. B. Distribution of cluster classifications across 100 repetitions for the comparison of dog and human β-CDR3. C. Distribution of cluster sizes across 100 repetitions for the comparison of dog and human β- CDR3. D. Comparison of the number of β-CDR3 sequences clustered or unclustered from dogs and humans across 100 repetitions. Each stacked bar represents one repetition. Blue, red, and green represent the counts of β-CDR3 sequences clustered from mice, both species, and humans, respectively. Light blue, orange, and light green bars represent the counts of unclustered β-CDR3 sequences from mice, both species, and humans, respectively. E. Distribution of cluster classifications across 100 repetitions for the comparison of mouse and human β-CDR3. F. Distribution of cluster sizes across 100 repetitions for the comparison of mouse and human β- CDR3.

## Discussion

The release of canine RNA-seq sequencing data and newly developed algorithms provide unprecedented opportunities to construct canine TCR repertoires. Although bulk RNA-seq provides essential insights into TCR repertoires, it also has certain inherent limitations. Bulk RNA-seq data is unable to catch the pair features of α-β chains within TCR. Additionally, the gene expression profiles obtained from bulk RNA-seq reflect the average gene expression of a cell population and is unable to decipher heterogeneities across different cell types. The single-cell (sc) RNA-seq data offer higher-resolution perceptions into gene expression at the cellular level. The scTCR-seq data allow us to investigate TCR light-heavy chain pairing and expression within individual cells. Integrating scRNA-seq and scTCR-seq data enables us to discern variations in TCR repertoires across distinct T-cell types.

Based on the current findings from new canine V gene and allele discovery, a significant portion of candidates remains elusive in the genome and lacks validation. Further validations are imperative. For instance, if the read distribution of variants in new gene and allele candidates closely resembles that of reference genes, we can reasonably conclude that the candidate is not a false positive. Another reason some candidates cannot be located in the genome is that certain new alleles are breed-specific. These alleles may only be expressed within a specific breed, yet no corresponding reference genome exists for them.

We have observed that the dominant DLA-I alleles vary among distinct dog breeds. We believe a similar phenomenon may occur in canine TCR germline repertoires. Therefore, studying the TCR repertoire within individual breeds and comparing it across different breeds is essential for gaining a deeper understanding of canine TCR repertoires.

We compared the diversities of dogs, mice, and humans. Due to the distinct abilities of different sequencing technologies to extract information from TCR repertoires, we consistently used the CDR3 obtained from TCR-seq for comparison. Notably, the mouse data we collected are from the same C3H strain, which could potentially explain the lower diversity observed in mouse CDR3.

## Material & Methods

### Data Collection

The canine RNA-seq dataset consists of 1,361 paired-end (PE) and 447 single-end (SE) samples from 44 PE and 17 SE BioProjects from the Sequence Read Archive (SRA) database. Due to the limited TCR identified in SE samples, we only focus on the TCR results for PE samples. The PE dataset includes 44 normal blood (BLO), 86 abnormal blood (ANB), 25 normal lymphoid tissue (NL), 38 abnormal lymphoid tissue (ANL), 15 T-cell lymphoma (TCL), 66 B-cell lymphoma (BCL), 488 tumor infiltrating T-cells (TIL), 320 normal tissue (NOR), 100 matched normal tissue (MNT), 129 non-cancer diseases (NCD), and 50 unclassified (UCF) samples. These samples were further classified into three categories: blood (BLO and ANB), lymphoid tissues (NL and ANL), and solid tissues (TIT, NOR, MNT, NCD and UCF). The top 5 cancers with biggest samples size in TIL dataset are 39 Glioma (GLI), 74 Hemangiosarcoma (HSA), 76 Oral melanoma (OM), 89 Osteosarcoma (OSA), and 161 Mammary tumors (MT) samples. The RNA-seq were collected from different batches, we conducted a comprehensive QC on RNA-seq data to ensure the data meet a series of quality standards and exclude failed samples to minimize potential artificial errors, as described in paper MHC-I landscape.

The human RNA-seq dataset comprises 1,231 bulk RNA-seq samples obtained from mammary tissue with breast cancer (BRCA) from TCGA^24^, along with 94 PE RNA-seq samples from blood with breast cancer from SRA^25^.

For the β- CDR3 comparison across dog, mice, and human, we used β-CDR3 obtained from TCR-seq. For canines, we gathered three bulk TCR-seq and 12 sc-TCR-seq samples. Bulk TCR-seq data for canines were processed using TRUST4 to reconstruct the TCR repertoires, while sc-TCR-seq data were analyzed using Cell Ranger to obtain TCR clonotypes. Mouse β-CDR3 sequences were sourced directly from a study that collected 12 mouse samples and conducted bulk TCR-seq^26^. Human β-CDR3 sequences were downloaded from TCRdb, a database containing human β-CDR3 data from TCR-seq experiments^14^.

The canine and human TCR germline repertoires were downloaded from IMGT. The canine genome assembly (canFam3.1, canFam4, canFam5, and canFam6) and gene annotation (canFam3 1.99 GTF) were obtained from the Ensembl database.

### TCR reconstruction

We used two software TRUST4^18^ and MiXCR^19^ to reconstruct canine TCR repertoires on all RNA-seq samples. We extracted complete CDR3 based on amino acid sequences which start with conserved cystine(C), end with Phenylalanine (F), no stop codons and reading frame shifting involved.

### Shuffling analysis to compare the quality of CDR3 from TRUST4 and MiXCR

Based on the intersection results in Figure 1A, we divided each type of CDR3 into three groups, TRUST4 only, overlap, and MiXCR only. To evaluate differences in length distributions between groups, we utilized the Jensen–Shannon divergence (JSD), with scores ranging from 0 to 1 indicating greater dissimilarity between distributions. We calculated the JSD for the length distribution of CDR3 in the overlap and MiXCR only groups. To avoid the errors induced by the imbalanced amount of CDR3 from distinct groups, firstly, we randomly sampled the same amount of CDR3 from TRUST4 unique as MiXCR unique. Second, we calculated the length distribution of the sampled TRUST4 only CDR3 and compared it with the length distribution of overlap CDR3, calculating the JSD score. Third, we compared the obtained JSD score with that of MiXCR unique and overlap CDR3. This process was repeated 10,000 times. Then, the p-value was computed using the equation: p = n/10,000, where n represents the frequency of instances where the JSD of overlap and sampled TRUST4 only CDR3 was lower than that of overlap and MiXCR only CDR3.

For motif conservation, we extracted the top three dominant length CDR3 sequences and conducted a similar shuffling analysis for each length. However, PCC was used to estimate motif similarity, with scores ranging from −1 to 1 indicating negative to positive correlation, respectively.

### Gene usage

We traced back to the TCR contigs corresponding to complete CDR3 and conducted further filtrations on contigs to obtain high-quality TCR contigs. Evaluation of TCR contigs focused on two criteria: overlap length and identity with reference genes. CDR3s corresponded to TCR contigs matching known V genes by more than 95%, with a matched length exceeding 50 nucleotides, were included in the gene usage analysis. This analysis was conducted for each respective category or group classification. Within each category, reads for each TRAV and TRBV gene were aggregated. To eliminate low-read V genes, a threshold of 50 total reads was established. Subsequently, the fraction of usage for each TRAV and TRBV gene was calculated, and gene usage was presented in descending order of frequency based on blood samples.

### Exploration of new variable genes and alleles in canine

We collected TCR contigs assembled by TRUST4 to explore the new TRAV and TRBV genes and alleles in canine. Each contig underwent alignment to known reference genes, and extraction was based on three nucleotide-level filtrations. First, the identity between contigs and known reference genes had to be less than 100%. Next, the alignment length between contigs and references for TRAV had to be greater than or equal to 245bp, or for TRBV, greater than or equal to 250bp. Third, the alignment should start at the first position of reference genes. Subsequently, two additional quality control measures at the amino acid level were applied. Contigs needed to translate to protein without reading frame shifts, and their protein sequences had to match conserved amino acids at specific positions such as 23C, 41W, and 104C. The contigs which passed the quality control measures were classified as new gene candidates and new allele candidates based on the identity score between contigs and known reference genes. Contigs with an identity score of less than 95% were classified as new gene candidates, while those with an identity score greater than or equal to 95% were classified as new allele candidates. However, some contigs exhibited high similarity to reference genes, with identities reaching 99% which might be caused by sequencing error. To avoid involving the contigs with sequencing errors, an additional filtration was conducted for contigs with less than 3 mismatches. If such contigs had a recurrence of greater than or equal to 2, they were classified as new allele candidates; otherwise, they were removed from the candidate list. Then, contigs were searched against three canine genomes (canFam4, canFam5, and canFam6), and those that could be 100% mapped to the genomes were retained. The RSS of those contigs were extracted from the genomes and compared to the RSS of reference genes. Contigs were classified as new gene or allele candidates if their RSS conservation matched that of the reference genes.

### CDR3 enrichment and depletion analysis

For each category and group classification of CDR3, we employed clusTCR to cluster the CDR3 sequences. Following clustering, we aggregated the reads supporting each CDR3 sequence. Subsequently, we focused on the top 10 largest clusters for downstream analysis. Utilizing the Fisher exact test, we analyzed the enrichment or depletion of CDR3 sequences among different tissue types based on their read counts. The p-values obtained from the Fisher exact test were adjusted using the Benjamini-Hochberg method and transformed to −log10. Enrichment or depletion was determined by assessing the odds ratio: if the odds ratio exceeded 1 and the associated q-value was greater than 0, it indicated enrichment; conversely, if the odds ratio was less than 1 and the q-value was negative, it suggested depletion.

### Comparison of **β**-CDR3 across dogs, mice, and humans

We gathered β- CDR3 from TCR-seq as described above. In total, we acquired 63,817 canine β- CDR3, 175,023 mouse β- CDR3, and 1,288,180 human β- CDR3. Due to the disparity in the number of CDR3s across species, we randomly sampled 63,817 β- CDR3 from mouse and human dataset 100 times separately to keep use the same amount of data to cluster. For each comparison between dog and human β-CDR3, we merged the canine and human sequences, removing any duplicates. We then applied clusTCR to cluster the β-CDR3 sequences, labeling the origin of each sequence accordingly. If a sequence was exclusively found in the dog dataset, it was labeled as “dog,” and the same process was repeated for human sequences. Sequences found in both datasets were labeled as “both.” Subsequently, we categorized the clusters into three groups: dog, both, and human. Clusters were labeled as “both” if more than 10% of the sequences originated from dogs and more than 10% from humans. Clusters where more than 90% of the sequences were from dogs were labeled as “dog,” and the same process was applied for human clusters. This analysis was repeated 100 times. The same process was followed for the comparison between mouse and human β-CDR3.

## Declarations

None

## Ethics approval and consent to participate

Not applicable.

## Consent for publication

Not applicable.

## Author contributions

All authors conducted the analysis, wrote, and approved the manuscript.

## Acknowledgments

We thank Dr. Yuan Feng for providing help on initiating this project; Ms. Jasmine Danielle Carter for editing the article; the Georgia Advance Computing Resource Center, University of Georgia (GACRC) for supporting this work. This work is funded by NCI R01 CA252713, R01 CA182093 and AKC Canine Health Foundation.

